# Rewarded extinction increases amygdalar connectivity and stabilizes long-term memory traces in the vmPFC

**DOI:** 10.1101/2021.12.08.471649

**Authors:** Nicole E. Keller, Augustin C. Hennings, Emily K. Leiker, Jarrod A. Lewis-Peacock, Joseph E. Dunsmoor

## Abstract

Neurobiological evidence in rodents indicates that threat extinction incorporates reward neurocircuitry. Consequently, incorporating reward associations with an extinction memory may be an effective strategy to persistently attenuate threat responses. Moreover, while there is considerable research on the short-term effects of extinction strategies in humans, the long-term effects of extinction are rarely considered. In a within-subjects fMRI study, we compared counterconditioning (a form of rewarded-extinction) to standard extinction, at recent (24 hours) and remote (∼1 month) retrieval tests. Relative to standard extinction, counterconditioning diminished 24-hour relapse of arousal and threat expectancy, and reduced activity in brain regions associated with the appraisal and expression of threat (e.g., thalamus, insula, periaqueductal gray). The retrieval of reward-associated extinction memory was accompanied by functional connectivity between the amygdala and the ventral striatum, whereas the retrieval of standard-extinction memories was associated with connectivity between the amygdala and ventromedial prefrontal cortex (vmPFC). One-month later, the retrieval of both standard- and rewarded-extinction was associated with amygdala-vmPFC connectivity. However, only rewarded extinction created a stable memory trace in the vmPFC, identified through overlapping multivariate patterns of fMRI activity from extinction to 24-hour and 1-month retrieval. These findings provide new evidence that reward may generate a more stable and enduring memory trace of attenuated threat in humans.

**Significance Statement:** Prevalent treatments for pathological fear and anxiety are based on the principles of Pavlovian extinction. Unfortunately, extinction forms weak memories that only temporarily inhibit the retrieval of threat associations. Thus, to increase the translational relevance of extinction research, it is critical to investigate whether extinction can be augmented to form a more enduring memory, especially after long intervals. Here, we used a multi-day fMRI paradigm in humans to compare the short- and long-term neurobehavioral effects of aversive-to-appetitive counterconditioning, a form of augmented extinction. Our results provide novel evidence that including an appetitive stimulus during extinction can reduce short-term threat relapse and stabilize the memory trace of extinction in the vmPFC, for at least one month after learning.

## Introduction

While learning about threats is adaptive, persistent and misattributed fearful responses are characteristic of anxiety disorders. Exposure therapy, based on the principles of Pavlovian extinction, is a widely used treatment for anxiety-related disorders (Abramowitz et al., 2019). Unfortunately, relapse of extinguished behavior is common, and a substantial number of individuals undergoing treatments will drop-out or relapse (Markowitz et al., 2015; Schottenbauer et al., 2008). Notably, even healthy adults tend to show post-extinction recovery of learned defensive behavior in new situations, indicating extinction is a fragile form of inhibitory learning bound to the spatiotemporal context in which extinction memories were formed (Bouton, 2002). Several augmented strategies to standard extinction have shown success in promoting relatively short-term (∼24 hours) retention of extinction memories in humans (Craske et al., 2018; Dunsmoor et al., 2015). However, evaluating the long-term success (> 1 week) of extinction protocols in humans is extremely rare, which limits the clinical translational relevance of extinction research, as symptoms frequently return some time after treatment (Vervliet et al., 2013). Here, we compared the neurobehavioral effects of standard extinction and augmented extinction in healthy adults at recent (24 hours) and remote (∼1 month) intervals in the same individuals. Whereas standard extinction involved simply omitting an expected aversive electrical shock, augmented extinction involved replacing the shock with a positive outcome, a paradigm known as aversive-to-appetitive counterconditioning (Dickinson & Pearce, 1977; Keller et al., 2020a).

In counterconditioning (CC), behavior is modified through a new association with a stimulus of the opposite valence. Research on counterconditioning dates to the earliest studies of conditioning in humans (Jones, 1924), and forms the basis for popular treatments for anxiety disorders such as systematic desensitization (Wolpe, 1954, 1968, 1995). Contemporary behavioral research on CC is sparse (Gatzounis et al., 2021; Keller & Dunsmoor, 2020; Koizumi et al., 2016; van Dis et al., 2019), and there are currently no neuroimaging studies directly comparing CC and extinction in humans. It remains unclear if a reduction of conditioned responses through CC is modulated by similar neural circuitry as standard extinction, and if the resulting threat attenuation is more enduring over time.

One possibility is that reduced relapse following CC is mediated by augmented activity in networks involved in the formation of extinction memories, specifically activity within and between the ventromedial prefrontal cortex (vmPFC) and amygdala (Giustino & Maren, 2015; Hartley & Phelps, 2010; Kredlow et al., 2021; Milad & Quirk, 2012). The presence of a positive stimulus could further engage reward-related regions of the mesostriatal dopamine system shown to be involved in threat extinction (Holtzman-Assif et al., 2010; Josselyn & Frankland, 2018; Kalisch et al., 2019; Luo et al., 2018; Raczka et al., 2011; Salinas-Hernández & Duvarci, 2021). In support of this idea, neurobiological evidence in rats found that rewarded extinction enhanced recruitment of an amygdala-striatal pathway and led to diminished threat relapse at a remote test (Correia et al., 2016). However, other evidence in rodents suggests that counterconditioning is less effective than standard extinction at preventing relapse of the original behavior (Holmes et al., 2016). If this were the case, then replacing shock with reward (rather than omitting it) may somehow interfere with processes underlying extinction memory formation and retrieval.

We developed a multi-day fMRI protocol to compare the neurobehavioral effects of threat extinction and aversive-to-appetitive CC on threat attenuation at recent and remote time points. The protocol incorporated a within-subjects Pavlovian conditioning design with renewal tests at 24-hours and approximately 1-month later. Based on our prior behavioral findings (Keller & Dunsmoor, 2020), we predicted CC would more effectively attenuate the relapse of conditioned responses. In line with previous research on enhanced extinction (Dunsmoor et al., 2019), we also predicted that CC would more effectively attenuate within-session activity in regions associated with threat appraisal (e.g., the insula, thalamus, dorsal anterior cingulate cortex (dACC), brainstem). Based on prior neurobiological evidence in rats (Correia et al., 2016), we also predicted that amygdala-ventral striatum functional connectivity would be selectively enhanced for stimuli associated with CC versus standard extinction.

To examine the fidelity of extinction and CC memory representations over time, we incorporated multivariate representational similarity analysis (Kriegeskorte et al., 2008) of encoding-to-retrieval overlap (Ritchey et al., 2012) between extinction learning and 24-hour and 1-month retrieval. We focused on the vmPFC based on recent fMRI evidence that 24-hour extinction retrieval reactivates similar neural activity patterns associated with extinction formation in this region (Hennings et al., 2020; Hennings et al., 2021). We predicted that neural similarity in the vmPFC would be enhanced and maintained over time for CC in comparison to standard extinction, indicating a more durable memory trace in a region critical for the encoding, storage, and retrieval of safety memories (Giustino & Maren, 2015; Milad & Quirk, 2012; Tovote et al., 2015).

## Materials and Methods

### Participants

Twenty-five participants (15 female; mean age: 23.48 years; SD = 5.51 years, age range 18-36), who reported no neurological or psychiatric disorders, were recruited from the University of Texas at Austin and local Austin community to complete this experiment. Two participants did not return for their third session ∼ 1 month away, therefore twenty-five participants completed the first and second session, and twenty-three (14 female; mean age 23.69 year; SD = 5.63 years, age range 18-36) completed all three sessions. We collected state individual difference measures [PTSD checklist for DSM-5 (PCL-5), the Childhood Trauma Questionnaire (CTQ), Beck Anxiety Inventory (BAI)] and trait measures [Intolerance of Uncertainty – Short Form (IUSF), State-Trait Anxiety Inventory (STAI)] of negative affect-related constructs for each participant. All participants provided written informed consent and procedures complied with the Institutional Review Board of UT Austin (IRB # 2017-02-0094).

### Task and Procedure

The design for this study was based on the behavioral experiment by (Keller & Dunsmoor, 2020). This was a within-subjects functional MRI study that included Pavlovian threat acquisition and extinction/counterconditioning on day one, and a renewal test and an episodic memory test 24 hours later and ∼ 1 month later (mean length: 26.91 days, SD = 10.26 days, day range: 15 to 63 days) depending on the participants’ availability (**Fig. 1A**). The episodic memory results are not discussed in this report. Conditioned stimuli (CS) were pictures of animals, tools, and food on a white background. The unconditioned stimulus (US) was a 6 msec electrical shock delivered to the index and middle finger of the participant’s left hand from the BIOPAC (Goleta, CA) MP160 System with a STM100C module. The task was presented using E-Prime 2.0 and consisted of a trial-unique category conditioning design, meaning that each trial was a different basic-level exemplar with a unique name. For example, there were not two different pictures of a dog. Pictures of common phobic stimuli (e.g., spiders, snakes, weapons) or highly appealing food items (e.g., pizza), were not used as CSs. For all phases (Pavlovian threat acquisition, extinction, and the renewal tests) each CS was on the screen for 6 seconds, followed by a 7-8 second ITI with a fixation cross on a white background. On each trial, subjects rated their expectancy to receive a shock using a two-alternative forced choice scale (2AFC) (i.e., yes or no). Trial order was pseudo-randomized so that participants did not see pictures from the same category three trials in a row.

**Figure 1.**
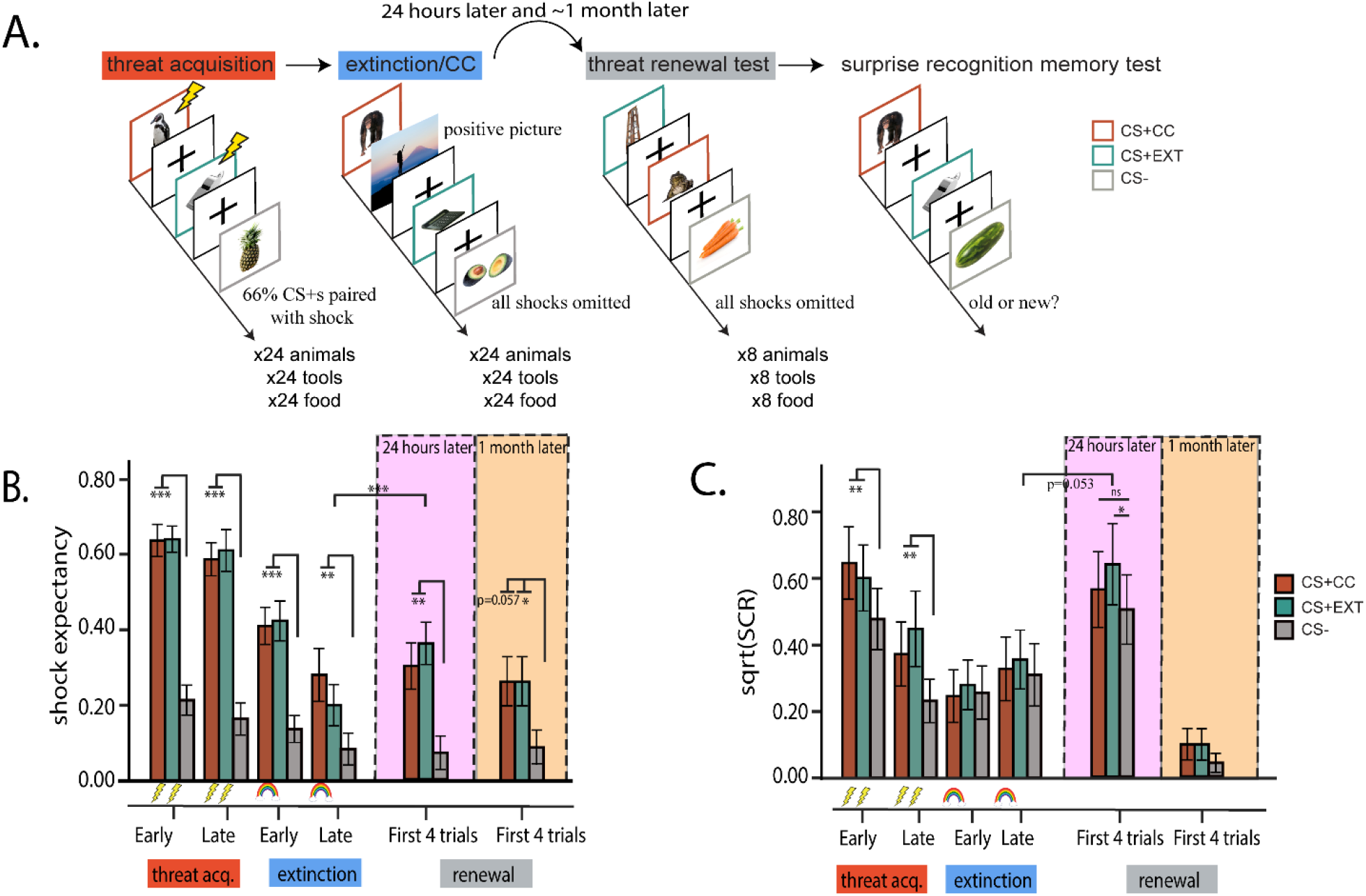
Experimental design and behavioral results. A.) Participants underwent threat acquisition with category exemplars of animals or tools (CS+s) paired with a shock on a partial reinforcement schedule, and a third category, food (CS-), never paired with shock. Conditioning was followed by the extinction phase, in which the shock was omitted following CS+EXT trials (counterbalanced, tools or animals, in this example tools), and CC, in which the shock was replaced by a positive picture at the end of each CS+CC trial (animals or tools, respectively, in this example animals). Subjects returned 24 hours and ∼1 month later, and new CSs from the same categories, were presented in the absence of any shocks or positive pictures. B.) Shock expectancy results confirmed successful acquisition and extinction of threat expectancy. 24 hours after day 1, shock expectancy towards the CS+EXT category significantly increased from the end of extinction to early renewal. Shock expectancy towards the CS+s remained even at the ∼1 month follow up. C.) Conditioned SCRs replicated prior findings (Keller & Dunsmoor, 2020), there were no differences between CS+s during acquisition or extinction, but 24 hours later, SCRs were higher for the CS+EXT category as compared to the CS-category, and there were no differences between the CS+CC category and the CS-category. 1 month later, there was no renewal of conditioned SCRs towards the CS+s. Colored borders are for illustrative purposed only. The rainbow, which represents the positive pictures, and the lightning bolt, which represents an electrical shock, depict the outcome following a given CS type. For example, following CS+CC trials during extinction, there is a positive picture. Error bars indicate SEM. P<0.001 (***), P<0.01 (**), P<0.05 (*).

#### Day 1

Day 1 included two phases: Pavlovian threat acquisition and extinction. These phases were divided into three separate functional imaging runs: 1) the first half of threat acquisition, 2) the second half of threat acquisition followed without a break by the first half of extinction, and 3) the last half of extinction. Each scanner run occurred consecutively with less than ∼ 1 min break between runs. Before participants entered the scanner, shock electrodes were attached to the index and middle finger on the left hand. The electrical shock was calibrated to be at a level that was deemed “highly annoying and unpleasant, but not painful”. Skin conductance response (SCR) served as a measure of conditioned autonomic arousal and was collected throughout the experiment on each day. SCR electrodes were placed on participants’ left palm and connected to a BIOPAC MP160 System (Goleta, CA). SCR sampling rate was set to 200Hz (see: Psychophysiology Analysis). Day 1 included a total of 144 trials across acquisition and extinction. During acquisition, CS stimuli from two categories (CS+, animals and tools, 24 stimuli per category) co-terminated with a shock 66% of the time. Items from the food category (CS-, 24 stimuli) were never paired to a shock, and always served as a within-subjects unpaired control category. Extinction included a total of 24 CS+CC, 24 CS+EXT and 24 CS-trials, all unpaired with shock. During extinction, stimuli from one CS+ category (CS+CC, animals or tools, counterbalanced across participants), were not followed by a shock and were paired with a unique image of a positive valenced picture, depicting a serene scene or accomplishment. The positive pictures were presented for a duration of 1 second and were pilot rated by a separate group of 19 participants to confirm high valence and low arousal. Stimuli from the other CS+ category (CS+ EXT, either tools or animals, respectively), were simply not followed by a shock.

#### Day 2

Participants returned the next day (∼24 hours) and underwent a test of threat renewal. The renewal test was followed by a recognition memory test for half the items encoded the previous day including the positive pictures paired with CS+CC during counterconditioning (details on the memory test and memory results not reported here). Before participants entered the scanner, shock and SCR electrodes were re-attached. Participants did not receive any new instructions from the day before and were instructed to continue to rate expectancy for receiving shocks on each trial. The renewal test included 8 trials each of animals, tools, and food. The CSs were novel category exemplars not shown the previous day. There were no shocks or positive pictures presented during the renewal test on Day 2. The first trial on the renewal test was always a discarded CS− trial that was used to capture the initial orienting response. Behavioral and neural analyses for the renewal test focused *a priori* on the first 4 trials (early renewal test) per CS type. Focusing on the first four trials is in line with previous human neuroimaging research on extinction recall (Kroes et al., 2016; Milad et al., 2009; Schiller et al., 2010), as these early trials capture the instance when the possibility for threat is most ambiguous. In the absence of a shock, later trials during the renewal test phase most likely reflect further extinction learning.

#### One month later

Participants returned for their third and final session ∼1 month later. This session followed the same format as Day 2, and included four functional imaging runs: a final threat renewal test, a recognition memory test for the rest of the CS exemplars from Day 1, and two runs of a perceptual category localizer. Before participants entered the scanner, shock and SCR electrodes were re-attached and the shock was re-calibrated.

### Psychophysiology Analysis

SCRs were calculated using prior criteria (Keller & Dunsmoor, 2020). SCRs were considered valid to the CS trial if the trough-to-peak deflection of electrodermal activity occurred between 0.5 to 6 seconds following CS onset and were not greater than 0.2 uS. Trials that did not meet these criteria were scored as zero. SCRs were scored by an automated analysis script implemented in Matlab (Green et al., 2014), and were later visually inspected by research assistants blind to the experimental conditions. SCR data were square-root transformed prior to statistical analysis to normalize the distributions. Participants were not excluded from the analysis based on any response criteria for SCRs, based on recommendations from the field of human threat conditioning (Lonsdorf et al., 2017). Two-AFC shock expectancy was coded as 1=expect to receive a shock, 0= do not expect.

### Imaging parameters

Brain images were recorded on a 3T Siemens Vida with 64-channel head coil at the University of Texas at Austin Biomedical Imaging Center. Functional task and localizer data were acquired using T2*-weighted EPI sequences (TR = 1000ms, TE = 86ms, FOV = 86 × 86mm, 2.5mm isotropic voxels), with slices oriented parallel to the hippocampal long axis and positioned to provide whole-brain coverage. High-resolution T1-weighted anatomical images were obtained using 3D MPRAGE sequences (TR = 2400ms, TE = 1000ms, FOV = 208 × 300mm, 0.8mm isotropic voxels) before the EPIs in each session, to aid in co-registration and normalization. Diffusion-weighted images were also acquired but were not examined.

### fMRI Data Preprocessing

MRI data were preprocessed using fMRIPrep 1.5.9 (Esteban et al., 2019) and FSL (FMRIB’s Software Library, www.fmrib.ox.ac.uk/fsl) FEAT 6.00 (FMRI Expert Analysis Tool). Processing in fMRIPrep followed the default steps, with additional options for multiple T1-weighted (T1w) images per participant (--*longitudinal* flag) and a framewise displacement threshold of 0.3mm. T1w images were corrected for intensity nonuniformity and skull-stripped using *N4BiasFieldCorrection* (Tustison et al., 2010) and *BrainExtraction* (both from ANTs 2.2.0; Avants et al. 2008). Segmentation of the skull-stripped T1w images into three tissue classes (CSF, WM, GM) was performed using FSL 5.0.9 *fast* (Y. Zhang et al., 2001), followed by surface reconstruction with FreeSurfer 6.0.1 *recon-all* (Dale et al., 1999). The skull-stripped T1w images were registered using FreeSurfer’s mri_robust_template to generate a single unbiased T1w-reference map per participant for spatial normalization (Reuter et al., 2010). Spatial normalization to MNI space was performed via nonlinear registration (ANTs *Registration*), using skull-stripped versions of both the T1w reference volume and MNI152NLin2009cAsym template (Fonov et al., 2009).

Functional data from each BOLD run were corrected for field distortion based on a B0-nonuniformity map estimated via AFNI *3dQwarp* (Cox & Hyde, 1997), then co-registered to the corresponding T1w reference using boundary-based registration (Greve & Fischl, 2009) with 6 degrees of freedom (FreeSurfer *bbregister*). Head-motion parameters, including transformation matrices and six rotation and translation parameters, were estimated for each BOLD run prior to any spatiotemporal filtering (FSL *mcflirt*). Framewise displacement and DVARS were calculated for each functional run using Nipype (Power et al., 2014), and frames exceeding 0.3mm FD or 1.5 standardized DVARS were annotated as motion outliers. In addition, six principal components of a combined CSF and white matter signal accounting for the most variance were extracted using aCompCor (Behzadi et al., 2007) following highpass filtering (128s cutoff) with discrete cosine filters. The BOLD runs were then slice-time corrected (AFNI *3dTshift*; Cox, 1996), and resampled onto original native space using custom methodology of fMRIPrep that applies all correction transformations in a single interpolation step. Additional details on the fMRIPrep pipeline may be found in the online documentation: https://fmriprep.org/en/1.5.9/.

Following preprocessing in fMRIPrep, we masked the preprocessed BOLD data for each participant with the intersection of the average T1-reference brain mask with the average BOLD reference mask. In final preparation of the MRI data for analysis with FSL (FMRIB’s Software Library, www.fmrib.ox.ac.uk/fsl, Version 6.00), the following pre-statistical processing was performed in FSL’s FEAT (FMRI Expert Analysis Tool): registration of the T1w-reference map and co-registration of the BOLD reference data to MNI152 space using *FLIRT* with 12 degrees of freedom (Jenkinson et al., 2002; Jenkinson & Smith, 2001), spatial smoothing using a Gaussian kernel of FWHM 5mm, and grand-mean intensity normalization of the entire 4D dataset by a single multiplicative factor.

Confound regressors consisting of the following MRIPrep-derived factors were prepared for functional denoising of individual BOLD runs: 6 aCompCor components, cosine filters for temporal filtering, 6 rotation and translation parameters and FD and spike regressors to exclude time points with excessive motion (>0.3mm FD or >1.5 standardized DVARS). MRIQC (Esteban et al., 2019) was used as a preliminary check of data quality. Scan runs were excluded from analysis if more than 20% of TRs exceeded a framewise displacement of 0.3mm. Only a single run (Functional run 2, Day 1) from one participant was excluded with this threshold.

### fMRI Analysis

fMRI analysis of the processed data was carried out using FEAT. Individual-level time-series statistical analyses were carried out using FILM with local autocorrelation correction (Woolrich et al., 2001). Separate regressors were specified for the experimental conditions of primary interest (CS+CC, CS+EXT, CS-) in each learning phase (threat acquisition: CS+s >CS-, CS->CS+s, extinction: CS+CC > CS+EXT, and renewal tests: CS+CC > CS+EXT), by convolving the stimulus function with a double-gamma hemodynamic response function (HRF), and adding a temporal derivative. Additional covariates included the electrical shock (following CS+ trials, during acquisition), positive pictures (following CS+CC trials, during counterconditioning), and confound regressors derived from fMRIPrep (described above). The higher-level analysis averaged contrasts estimates in each learning phase (acquisition, extinction/CC and the renewal tests), and was carried out using FLAME (FMRIB’s Local Analysis of Mixed Effects) stage 1 (Beckmann et al., 2003; Woolrich, 2008; Woolrich et al., 2004). Whole-brain Z (Gaussianised T/F) statistic images were thresholded non-parametrically using clusters determined by Z>3.1 and a (corrected) cluster significance threshold of P=0.05 (Worsley, 2001). A left superficial amygdala mask from the Juelich histological atlas (Amunts et al., 2005; Eickhoff et al., 2005), with a probability threshold of 30%, was used as a pre-thresholding mask for analysis of 24-hour renewal. We then performed a small-volume correction (SVC) within this mask identified at Z>3.1 and cluster corrected at p=0.05 (Worsley, 2001). Anatomical labels in the tables of activation were obtained by converting significant cluster coordinates in MNI space to Talairach space using GingerALE 3.0.2 (www.brainmap.org) (Laird et al., 2010), and subsequently using Talairach Client (Lancaster et al., 2000).

### Region of Interest selection

*A priori* ROIs for parameter estimate analysis included brain regions that are reliably characterized in meta-analyses of Pavlovian threat conditioning and extinction studies, and are involved in threat expression: the insula, dACC, thalamus and brainstem (periaqueductal gray) (Fullana et al., 2016; Fullana et al., 2018). For each of these threat ROIs, a sphere was drawn around peak coordinates reported in these studies, with a radius of 10mm. Parameter estimates for ROIs were extracted using FSL’s featquery tool and input to R Studio for further analyses with paired *t*-tests. The vmPFC, an *a priori* ROI for functional connectivity and RSA analyses, was defined functionally from the CS-> CS+s contrast during acquisition. A 10 mm sphere was drawn around the coordinates of a significant cluster (z<3.1, cluster corrected p<0.05) corresponding to the medial frontal gyrus (MNI coordinates, -14, 50, -1) (Table 1).

**Table 1.** Single group average (paired t-test) whole-brain contrasts during threat acquisition, identified at Z > 3.1 (cluster-corrected p < 0.05)

| Contrast | Region | MNI coordinates |  |  | Size (voxels) | Cluster P | Cluster Z |
| --- | --- | --- | --- | --- | --- | --- | --- |
|  |  | x | y | z |  |  |  |
| CS+s > CS- |  |  |  |  |  |  |  |
|  | Inferior Occipital Gyrus, BA 19 | -46 | -75 | 2 | 13676 | 3.50E-19 | 7.02 |
|  | Inferior Temporal Gyrus, BA 37 | 51 | -71 | 2 | 11861 | 2.21E-17 | 7.96 |
|  | Cuneus, BA 18 | 7 | -94 | 20 | 3206 | 7.15E-07 | 5.32 |
|  | Precuneus, BA 7 | -8 | -70 | 46 | 2351 | 1.90E-05 | 4.58 |
|  | Superior Frontal Gyrus, BA 6 | 5 | 8 | 70 | 1541 | 0.000612 | 4.72 |
|  | Posterior Cingulate, BA 23 | -1 | -27 | 22 | 952 | 0.0114 | 4.56 |
|  | Inferior Frontal Gyrus, BA 47 | -33 | 23 | -10 | 926 | 0.0132 | 4.27 |
| CS- > CS+s |  |  |  |  |  |  |  |
|  | Postcentral Gyrus, BA 3 | -13 | -33 | 74 | 24383 | 1.56E-28 | 6.04 |
|  | Clastrum | -35 | -9 | 9 | 5725 | 1.99E-10 | 5.06 |
|  | Lingual Gyrus, BA 18 | 20 | -82 | -8 | 4205 | 2.33E-08 | 5.38 |
|  | Clastrum | 38 | -4 | 4 | 3305 | 5.36E-07 | 6.07 |
|  | Lingual Gyrus. BA 8 | -17 | -94 | -9 | 3235 | 6.56E-07 | 5.44 |
|  | Medial Frontal Gyrus, BA 10 | -14 | 50 | -1 | 2516 | 9.89E-06 | 4.4 |
|  | Superior Temporal Gyrus, BA 41 | 58 | -17 | 6 | 1493 | 0.000765 | 4.13 |
|  | Lingual Gyrus, BA 18 | -20 | -58 | 8 | 1373 | 0.00135 | 5.15 |
|  | Parahippocampal Gyrus, Hippocampus | -28 | -15 | -24 | 1072 | 0.00606 | 4.79 |
|  | Posterior Cingulate, BA 30 | 22 | -53 | 8 | 1019 | 0.008 | 3.92 |
| CS+CC > CS+EXT |  |  |  |  |  |  |  |
|  | Postcentral Gyrus, BA 3 | -44 | -25 | 55 | 912 | 0.0188 | 3.86 |
|  | Middle Temporal Gyrus, BA 37 | -59 | -60 | 0 | 825 | 0.0297 | 4.56 |
| CS+EXT>CS+CC | No significant activity | — | — | — | — | — | — |

### Task-Based Functional Connectivity

We used generalized psychophysiological interaction (gPPI) to examine functional connectivity at the 24-hour and ∼1-month renewal tests, in two *a priori* pathways (basolateral amygdala (BLA) →nucleus accumbens (NAc) and vmPFC→central amygdala (CeM)). The timeseries for the seeds (BLA and vmPFC) were extracted using FSL’s meants command and input as regressors in the model. Interactions between the physiological variable (i.e., the seed’s respective timeseries) and each of the psychological variables (i.e., CS+CC, CS+EXT and CS-) were computed and included in the design matrix as the variables of interest. In accordance with human neuroimaging research on the circuitry of amygdala subregions (Koch et al., 2016; Roy et al., 2009, 2013), standardized amygdala ROIs (BLA and CeM) were defined using the Juelich histological atlas (Amunts et al., 2005; Eickhoff et al., 2005) as implemented in FSL. Following (Koch et al., 2016), voxels were included if they had a 50% or higher probability of belonging to the CeM, but due to signal drop-out in the temporal cortex, we used a more stringent threshold of 70% for the BLA. The anatomically defined NAc seed was derived from the Harvard-Oxford Subcortical Probability Atlas, thresholded at 50%. The vmPFC was defined functionally from the CS-> CS+ contrast during acquisition (see: Region of Interest selection).

Mean z-scores of connectivity from target ROIs were extracted using Featquery for each regressor of interest (CS+CC, CS+EXT and CS-), at both the 24-hour and ∼1 month renewal tests. These connectivity means were then input into R studio for further statistical analyses.

#### Representational Similarity Analysis (RSA)

In order to facilitate RSA, LS-S style betaseries were computed for each scanner run (Mumford et al., 2012, 2014). Within each scanner run trial-specific beta images were iteratively computed in FEAT using a design matrix which modeled a single trial of interest and all of trials as regressors of no interest based on trial type (e.g., separate CS+CC, CS+EXT, CS-regressors of no interest). FEAT settings were identical as in our univariate analysis, with the exception that no spatial smoothing was applied in order to respect the boundaries of our *a priori* ROIs in multivariate analyses. In addition to these trial-specific beta estimates, we also generated conventional estimates of average activity for each CS type during each phase (i.e., all CS+CC in one regressor of interest), again without spatial smoothing. For the renewal sessions, separate regressors were used to model the early vs. late trials.

RSA was accomplished using custom Python code. The goal of our analyses was to iteratively compare multivoxel patterns of activity in the vmPFC, between memory encoding in the extinction/CC session, recent renewal, and remote renewal. In order to reduce noise across the multivoxel pattern prior to estimating pattern similarity, each LS-S beta image was weighted (multiplied) by the overall univariate activity estimate of the corresponding CS type and time point (Hennings et al., 2020; Hennings et al., 2021; H. Kim et al., 2020) (e.g., all images of CS+CC from early 24-hour renewal were weighted with the average CS+CC pattern from the same time point). For each CS type, all of the LS-S images were entered into a representational similarity matrix, where each cell represents the Pearson’s correlation of the multivoxel patterns of activity between two images in the vmPFC. For each CS type, the correlations were fisher-z transformed, and the average similarity was taken for our three comparisons of interest: extinction/CC encoding to recent renewal, extinction/CC encoding to remote renewal, and recent renewal to remote renewal. Average fisher-z similarity values were then exported to R studio for statistical analysis.

### Analytic Plan

All statistical analyses were carried out in the R environment (Team, 2013). Data was analyzed using repeated measures analysis of variance (ANOVA), with the *ez* package (Lawrence, 2016), and included factors for CS Type (CS+CC,CS+EXT, and CS-) and time (e.g., first and second half of phase, or recent and remote renewal phases) where appropriate. Greenhouse-Geisser (GG) correction was applied when sphericity was violated. Main effects or interactions were followed by *post-hoc* two-tailed paired *t-tests*.

## Results

### Behavioral Results

#### Threat acquisition and extinction

Analyses of mean shock expectancy and SCRs during the acquisition and extinction phases on Day 1 were separated into the first and second half of trials (i.e., early/late) (**Fig. 1B, 1C**). Shock expectancy was significantly higher for both CS+s in comparison to CS-during both early and late trials of acquisition (all *p <* 0.001) (**Fig. 1B**). A repeated-measures ANOVA of SCR during acquisition revealed a main effect of CS type (F_(1.50, 36.04)_ = 11.462, *p*_*gg*_ < 0.001, η^2^_G_ = 0.025) and a main effect of early/late trials (F_(1,24)_ = 21.194, *p* < 0.001, η^2^_G_ = 0.053), but no interaction (*p*_*gg*_ = 0.071). *Post-hoc* paired *t*-tests showed successful acquisition towards both CS+s, as SCRs were significantly higher for CS+CC vs CS- and CS+EXT vs CS-(all *p* < 0.01) (**Fig. 1C)**. Importantly, shock expectancy and SCR did not differ between CS+s during acquisition. Thus, participants successfully acquired equivalent expectancy responses and conditioned arousal towards both CS+s.

A repeated measures ANOVA of shock expectancy during extinction revealed a significant main effect of CS Type (F_(1.89, 45.42)_ = 12.810, *p*_*gg*_ < 0.001, η^2^_G_ = 0.143), early/late trials (F_(1, 24)_ = 10.440, *p* = 0.004, η^2^_G_ = 0.066) and an interaction of CS Type by early/late trials (F = 5.514, *p*_gg_ = 0.013, η^2^_G_ = 0.018). While mean shock expectancy ratings were still significantly higher for CS+s in comparison to CS-during the first half (all *p* < 0.001), and second half (all *p* ≤ 0.01) of extinction, there was a significant decrease in shock expectancy for CS+EXT stimuli from the first to the last half of extinction (t_(24)_ = 5.073, *p* < 0.001, 95% CI [0.133, 0.314]), but not for CS+CC stimuli (*p* = 0.069) (**Fig. 1B**). A repeated-measures ANOVA of SCR means from extinction showed no effect of CS Type (*p*_*gg*_ = 0.471), no effect of early/late trials (*p* = 0.237), nor an interaction (*p*_*gg*_ = 0.786), indicating successful diminishment of conditioned SCRs via the absence of shock (**Fig. 1C**).

#### 24-hour threat renewal test

Mean shock expectancy during early 24-hour renewal (first 4 trials) was higher for both CS+s in comparison to CS− (all *p* <0.01), and there were no differences between CS+s (*p* = 0.387) (**Fig. 1B**).

Notably, given the limited sensitivity of a 2AFC, we did not expect to see differences between CS+s within sessions. As such, we assessed expectancy during the end of extinction, and compared it to expectancy during the renewal phase. A repeated measures ANOVA with a factor of CS Type and phase (last half of extinction and early renewal), revealed a main effect of CS Type (F_(1.73,41.56)_ = 11.26, *p*_*gg*_ < 0.001, η^2^_G_ = 0.115), a trend toward a significant main effect of phase (F_(1,24)_ = 4.04, *p* = 0.056, η^2^_G_ = 0.010), but no significant CS Type by phase interaction (*p*_*gg*_ = 0.072). *Post-hoc* paired *t*-tests revealed that expectancy for CS+EXT significantly increased (t_(24)_ = 3.894, *p* < 0.001, 95% CI [0.075, 0.245]) from late extinction to early renewal, but was not different between phases for neither CS+CC (*p* = 0.720) nor CS-stimuli (*p* = 0.818). Thus, at 24 hours, participants exhibited renewal of shock expectancy towards items from the category that underwent standard extinction, but not towards items from the control category, nor the CC category.

Repeated-measures ANOVA of SCRs during 24-hour renewal revealed a main effect of CS type (F_(1.81,43.39)_ = 3.732, *p*_*gg*_ = 0.036, η^2^_G_ = 0.010) (**Fig. 1C**). *Post-hoc* paired *t*-tests revealed greater mean SCRs towards CS+EXT versus CS− (t_(24)_ = 2.374, *p* = 0.026, 95% CI [0.018, 0.255]), but no difference between CS+CC versus CS− (*p* = 0.186), nor CS+CC versus CS+EXT (*p* = 0.122). Thus, while SCRs did not differ between CS+s, participants expressed heightened conditioned arousal to items from the CS+EXT category as compared to items from the CS-category, but this difference was eliminated for CS+CC stimuli.

An ANOVA comparing physiological arousal at the end of extinction to early renewal revealed a main effect of CS Type (F_(1.90,45.54)_ = 4.099, *p*_*gg*_ = 0.025, η^2^_G_ = 0.005), no main effect of phase (*p* =0.062) and no significant CS Type by phase interaction (*p*_*gg*_ = 0.354). *Post-hoc* paired *t*-tests revealed that conditioned arousal for CS+EXT stimuli was marginally higher (t_(24)_ = 2.037, *p* = 0.053, 95% CI [-0.004, 0.5693]) from late extinction to early renewal, but was not different between phases for neither CS+CC (*p*= 0.081) nor CS-(*p*=0.089) stimuli.

### 1 month threat renewal test

Approximately 1 month later, participants did maintain slightly elevated shock expectancy to each CS+ versus the CS-(**Fig. 1B**). While a repeated measures ANOVA of mean shock expectancy revealed no significant main effect of CS Type (*p*_*gg*_ = 0.080), *post-hoc* paired *t*-tests revealed significantly higher expectancy for CS+EXT in comparison to the CS-(t_(24)_ = 2.336, *p* = 0.029, 95% CI [0.0195, 0.328]), a trend towards significantly higher shock expectancy for CS+CC in comparison to the CS-(t_(24)_ = 2.005, *p* = 0.057, 95% CI [-0.001, 0.354]), and no differences between CS+s (*p* = 1). Interestingly, autonomic arousal to each CS was exceptionally low (**Fig. 1C**). A repeated measures ANOVA of mean SCR revealed no main effect of CS type (*p*_*gg*_ = 0.395). Thus, 1 month later, participants expressed some retrieval of Day 1 CS+ shock contingencies, but did not display heightened physiological arousal towards CS+ items.

## Neuroimaging Results

### Univariate analysis

#### Extinction

Univariate whole-brain fMRI analysis focused on the extinction and renewal test phases (see Tables 1-4 for full results from each experimental phases). During extinction, a contrast of CS+CC > CS+EXT revealed significant clusters only in the cuneus and precuneus (**Table 2**). The inverse contrast (CS+EXT > CS+CC) revealed significant clusters in brain regions traditionally involved in maintaining and expressing threat (Fullana et al., 2016) (**Table 2, Fig. 2A**). To further characterize these fMRI results, we extracted activity associated with each stimulus type (CS+CC, CS+EXT and CS-) from *a priori* regions of interest putatively involved in acquisition and extinction of threat (Fullana et al., 2016; Fullana et al., 2018) (i.e., dACC, insula, thalamus and PAG). We focused these ROI analyses on the second half of extinction. This revealed diminished activity to the CS+CC in comparison to the CS+EXT (**Fig. 2C**), indicating that the outcome during counterconditioning attenuated activity in regions involved in maintaining and expressing threat expectations relative to merely omitting the shock.

**Table 2.** Single group average (paired t-test) whole-brain contrasts during extinction identified at Z > 3.1 (cluster-corrected p < 0.05)

| Contrast | Region | MNI coordinates |  |  | Size (voxels) | Cluster P | Cluster Z |
| --- | --- | --- | --- | --- | --- | --- | --- |
|  |  | x | y | z |  |  |  |
| **CS+s>CS-** |  |  |  |  |  |  |  |
|  | Superior Frontal Gyrus, BA 6 | -2 | 25 | 62 | 3335 | 5.36E-07 | 5.06 |
|  | Inferior Temporal Gyrus, BA 37 | 52 | -70 | 0 | 1682 | 0.000352 | 4.77 |
|  | Precuneus, BA 7 | 2 | -74 | 50 | 1216 | 0.00312 | 4.87 |
| **CS->CS+s** |  |  |  |  |  |  |  |
|  | Declive | 19 | -76 | -10 | 59754 | 0 | 6.99 |
|  | Medial Frontal Gyrus, BA 6 | -4 | -5 | 55 | 39827 | 1.75E-39 | 5.79 |
|  | Insula, BA 13 | 36 | -26 | 16 | 13300 | 1.20E-18 | 6.2 |
|  | Insula, BA 13 | -37 | -26 | 15 | 6292 | 4.77E-11 | 5.11 |
|  | Inferior Occipital Gyrus, BA 17 | -13 | -94 | -7 | 4245 | 2.44E-08 | 5.38 |
|  | Precentral Gyrus, BA 4 | 55 | -12 | 37 | 2726 | 4.95E-06 | 4.36 |
|  | Precentral Gyrus, BA 6 | -44 | 0 | 30 | 1786 | 0.000223 | 4.62 |
|  | Middle Temporal Gyrus, BA 19 | 38 | -63 | 19 | 1306 | 0.00201 | 4.47 |
|  | Precuneus, BA 7 | 24 | -54 | 47 | 1231 | 0.0029 | 4.36 |
|  | Anterior Cingulate, BA 32 | 8 | 50 | -9 | 1065 | 0.00669 | 4.08 |
|  | Precentral Gyrus, BA 6 | 45 | 3 | 31 | 962 | 0.0115 | 4.28 |
|  | Inferior Frontal Gyrus, BA 47 | 28 | 31 | -11 | 738 | 0.0396 | 4.6 |
|  | Superior Temporal Gyrus, BA 22 | -51 | 1 | -9 | 714 | 0.0454 | 3.74 |
| **CS+ CC > CS+EXT** |  |  |  |  |  |  |  |
|  | Precuneus, BA 7 | 4 | -65 | 59 | 1949 | 5.10E-05 | 6.3 |
|  | Cuneus, BA 19 | 3 | -85 | 36 | 680 | 0.04 | 4.56 |
| **CS+EXT>CS+CC** |  |  |  |  |  |  |  |
|  | Culmen | 18 | -37 | -13 | 117657 | 0 | 6.68 |
|  | Postcentral Gyrus, BA 2 | -36 | -24 | 45 | 12787 | 1.95E-19 | 5.84 |
|  | Inferior Frontal Gyrus, BA 9 | 47 | 14 | 26 | 6756 | 1.90E-12 | 4.4 |
|  | Cingulate Gyrus, BA 24 | -6 | 4 | 49 | 6719 | 2.12E-12 | 5.34 |
|  | Clastrum | 32 | 24 | -1 | 2292 | 1.12E-05 | 5.68 |
|  | Precuneus, BA 7 | 28 | -52 | 45 | 1722 | 0.000146 | 5.19 |
|  | Superior Temporal Gyrus, BA 22 | 61 | -39 | 14 | 1371 | 0.000814 | 4.48 |
|  | Precentral Gyrus, BA 4 | 46 | -15 | 36 | 993 | 0.00617 | 4.15 |
|  | Superior Temporal Gyrus, BA 38 | 50 | 9 | -19 | 791 | 0.0201 | 5.04 |

**Figure 2.**
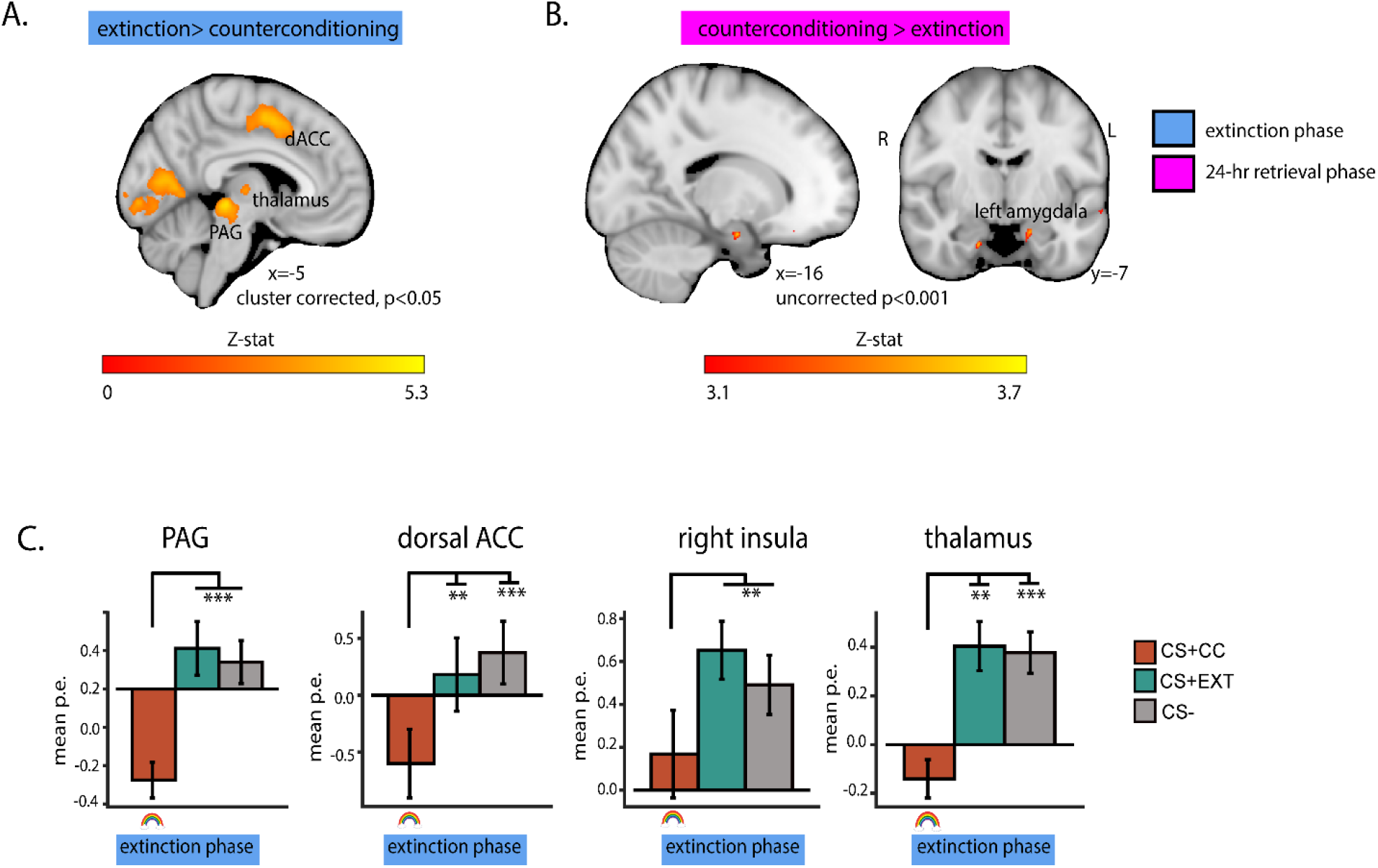
CC was associated with reduced activity in threat ROIs during extinction, and enhanced amygdala activation during 24-hour renewal. A.) A whole-brain contrast of CS+EXT > CS+CC during the extinction phase, identified at Z>3.1, cluster corrected at p<0.05, revealed activity in regions traditionally associated with threat appraisal and expression (e.g., periaqueductal grey, dACC, insula and thalamus). B.) A whole-brain contrast of CS+CC > CS+EXT during the 24-hour renewal phase, identified at p<0.001 uncorrected for multiple comparisons, revealed a cluster in the left amygdala. C.) Parameter estimates extracted from a priori regions associated with threat (periaqueductal grey, dACC, insula and thalamus), during the last half of extinction, revealed significantly lower activity for CS+CC stimuli in comparison to both CS+EXT and CS-stimuli. The rainbow, which represents the positive pictures, depict the outcome following CS+CC stimuli. Error bars indicate SEM. P<0.001 (***), P<0.01 (**), P<0.05 (*).

#### 24-hour threat renewal test

Univariate fMRI analysis of the CS+EXT > CS+CC and CS+CC > CS+EXT contrasts did not reveal any significant activity that survived whole-brain correction for multiple comparisons. A more liberal exploratory threshold of *p* < 0.001 (uncorrected) for the CS+CC > CS+EXT contrast revealed a cluster in the left amygdala (MNI -16, -7, -21; 27 voxels, z = 3.49, *p*_*uncorrected*_ < 0.001; cluster corrected at *p* < 0.05 with SVC) (**Table 3, Fig. 2B**). No regions emerged at this liberal threshold for the inverse contrast (CS+EXT > CS+CC).

**Table 3.** Single group average (paired t-test) whole-brain contrasts during 24-hour threat renewal, identified at Z > 3.1 (cluster-corrected p < 0.05)

| MNI coordinates |  |  |  |  |  |  |  |
| --- | --- | --- | --- | --- | --- | --- | --- |
| Contrast | Region | x | y | z | Size (voxels) | Cluster P | Cluster Z |
| **CS+s > CS-** | No significant activity | — | — | — | — | — | — |
| **CS- > CS+s** | No significant activity | — | — | — | — | — | — |
| **CS+CC > CS+EXT (SVC of left amygdala, cluster corrected p<0.05)** | Left amygdala | -16 | -7 | -21 | 27 | 0.0185 | 3.49 |
| **CS+EXT > CS+CC** | No significant activity | — | — | — | — | — | — |

**Table 4.** Single group average (paired t-test) whole-brain contrasts during ∼1 month fear retrieval, identified at Z > 3.1 (cluster-corrected p < 0.05)

| **Contrast** | **Region** | **x** | **y** | **z** | **Size (voxels)** | **Cluster P** | **Cluster Z** |
| --- | --- | --- | --- | --- | --- | --- | --- |
| **CS+s > CS-** | No significant activity | — | — | — | — | — | — |
| **CS- > CS+s** |  |  |  |  |  |  |  |
|  | Precentral Gyrus, BA 4 | 48 | -12 | 42 | 3335 | 1.90E-08 | 5.06 |
|  | Paracentral Lobule, BA 3 | 21 | -32 | 58 | 1682 | 2.58E-08 | 4.77 |
|  | Declive | 18 | -81 | -13 | 1216 | 0.01 | 4.87 |
| **CS+CC > CS+EXT** | No significant activity | — | — | — | — | — | — |
| **CS+EXT > CS+CC** | No significant activity | — | — | — | — | — | — |

#### 1 month threat renewal test

No regions emerged at the whole-brain level for the univariate contrasts CS+CC > CS+EXT or CS+EXT > CS+CC at 1 month, even using a liberal threshold (*p* < .001, uncorrected).

### Functional connectivity

#### A BLA→NAc circuit for retrieval of rewarded extinction

To examine the involvement of fMRI derived amygdala projections, we conducted a generalized psychophysiological interaction analysis (gPPI) during recent and remote threat renewal tests (**Fig. 3A**). This analysis was inspired by neurobiological evidence that a BLA to NAc circuit preferentially supports reduced threat relapse of rewarded extinction (Correia et al., 2016). The seed region was an anatomically defined BLA, and the target region was an anatomically defined NAc.

**Figure 3.**
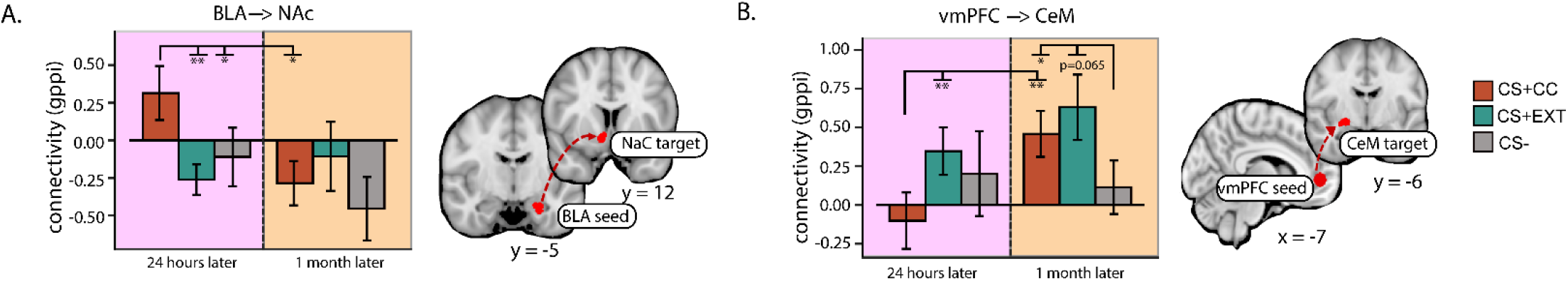
Functional connectivity during recent and remote renewal tests in two a priori pathways. A.) Functional connectivity using the BLA as a seed region and the NAc as a target region, was enhanced for CS+CC stimuli during 24-hour renewal, but was not different between CS types at ∼1 month renewal. B.) Functional connectivity using the vmPFC as a seed region and the CeM as a target region, was enhanced for CS+EXT, in comparison to CS+CC stimuli during 24-hour renewal. At ∼1 month, connectivity for CS+CC stimuli significantly increased, and was not different than CS+EXT stimuli. At this remote timepoint, both CS+s were associated with a greater functional vmPFC→CeM connection than CS-stimuli. Error bars indicate SEM. P<0.001 (***), P<0.01 (**), P<0.05 (*).

Twenty-four hours following extinction, a repeated measures ANOVA revealed a main effect of CS Type (F_(1.93, 46.24)_ = 5.781, *p*_*gg*_ = 0.006, η^2^_G_ = 0.085). *Post-hoc* paired *t*-tests revealed that connectivity between the BLA and the NAc, at this recent timepoint, was enhanced for stimuli from the CS+CC category, in comparison to stimuli from the CS+EXT category (t_(24)_ = 3.320, *p* = 0.003, 95% CI [0.217, 0.932]) and the CS-category (t_(24)_ = 2.631, *p* = 0.015, 95% CI [0.091, 0.756]). One month after extinction, there were no differences in connectivity between CS types (all p > 0.3). But, comparing across renewal test intervals (recent versus remote), *post-hoc* paired *t*-tests revealed that BLA→NAc connectivity significantly diminished for CS+CC stimuli from the 24-hour to the ∼1-month renewal test (t_(22)_ = -2.087, *p* = 0.048, 95% CI [-0.990, -0.003]).

#### A vmPFC→CeM circuit is recruited for CS+ stimuli at a remote renewal test

The medial PFC is considered a critical region that inhibits conditioned defensive responses via projections that inhibit the central nucleus of the amygdala (CeM) (Ghashghaei & Barbas, 2002; McDonald et al., 1996).This circuit is considered critical for successful extinction retrieval. We therefore conducted a gPPI during recent and remote threat renewal tests using the vmPFC as the seed region and an anatomically defined region of the CeM as the target region (**Fig. 3B**). The vmPFC was functionally defined based on a medial frontal gyrus cluster from the CS-> CS+ contrast during acquisition (**Table 1**), as anatomical labels for the vmPFC are variable across studies of Pavlovian conditioning and extinction.

Twenty-four hours following extinction, *post-hoc* paired *t*-tests revealed that connectivity between the vmPFC→CeM was heightened for CS+EXT stimuli versus CS+CC (t_(24)_ = 2.999, *p* = 0.006, 95% CI [0.140, 0.755]). One month following extinction, *post-hoc* paired *t*-tests revealed that connectivity between CS+CC and CS+EXT stimuli no longer differed (*p =* 0.466). At this remote timepoint, CS+CC stimuli (t_(22)_ = 2.250, *p* = 0.035, 95% CI [0.027, 0.661]) showed stronger vmPFC→CeM connectivity than the CS-stimuli, but there were no differences between CS+EXT and CS-stimuli (*p* = 0.065). Finally, there was a significant main effect of CS Type (F_(1.75, 38.49)_ = 5.93, *p*_*gg*_ = 0.008, η^2^_G_ = 0.043), and renewal test interval (F_(1, 22)_ = 5.24, *p =* 0.032, η^2^_G_ = 0.047), but no significant CS Type by renewal interval interaction (*p*_*gg*_ = 0.328). *Post-hoc* paired *t*-tests revealed that vmPFC→CeM connectivity for CS+CC significantly increased from the 24-hour to the 1-month renewal test (t_(22)_ = 3.370, *p =* 0.003, 95% CI [0.239, 1.005]).

### Multivariate representational similarity analysis

#### Pattern similarity between extinction/CC memory encoding retrieval

To assess the fidelity of the extinction and CC memory traces over time, we used RSA (Kriegeskorte et al., 2008) to compare patterns of fMRI activity during extinction/counterconditioning and 24-hour and 1-month renewal tests. We focused this analysis on the vmPFC, as this region is associated with successful extinction recall in humans (Milad et al., 2007; Phelps et al., 2004). Voxel-wise patterns of activity elicited by CS+CC, CS+EXT and CS-stimuli, were correlated with the pattern of activity elicited by novel stimuli from the same categories at the renewal test 24 hours (extinction → 24-hour renewal), ∼1 month later (extinction → 1 month renewal), and across renewal sessions (24-hour renewal → 1 month renewal). Notably, one innovation to the category-conditioning design (Dunsmoor et al., 2014; Hennings et al., 2020) is that participants are exposed to new category exemplars composing each CS category. Thus, pattern similarity cannot be driven simply by perceptual overlap of CSs, as different basic level items are presented at each phase.

#### A CC memory trace is stable in the vmPFC from encoding to recent and remote renewal tests

A repeated measures ANOVA on pattern similarity from encoding to recent renewal (extinction→24-hour renewal), revealed a main effect of CS Type (F_(1.75, 41.88)_ = 4.20, *p*_*gg*_= 0.026, η^2^_G_ = 0.081) (**Fig. 4A**). *Post-hoc* paired *t*-tests revealed that at 24 hours similarity from encoding to retrieval in the vmPFC was selectively enhanced for CC stimuli in comparison to CS+EXT stimuli (t_(24)_ = 2.169, *p* = 0.040, 95% CI [0.003, 0.110]) and CS-stimuli (t_(24)_ = 2.491, *p* = 0.020, 95% CI [0.001, 0.105]). At ∼1 month (extinction → 1 month renewal), neural similarity for CS+CC stimuli was enhanced in comparison to CS-stimuli (t_(22)_ = 2.147, *p*_*gg*_ = 0.043, 95% CI [0.002, 0.093]) (**Fig. 4B)**. Notably, memory traces from the extinction phase on Day 1 did not significantly change from recent to remote renewal, as a repeated measures ANOVA with factors of CS Type and renewal phase (extinction→24 hour renewal and extinction → 1 month renewal) revealed no main effect of phase (*p*_*gg*_ = 0.286), a significant main effect of CS Type (F_(1.61, 35.35)_ = 5.73, *p*_*gg*_ = 0.011, η^2^_G_ = 0.077), but no CS Type by phase interaction (*p*_gg_ = 0.702). **(Fig. 4A, 4B)** Thus, both at recent and remote timepoints, the CC memory trace was stable in the vmPFC.

#### Similarity patterns in the vmPFC across recent and remote renewal are enhanced for CC stimuli

A repeated measures ANOVA of pattern similarity from recent to remote renewal (24-hour renewal session→ 1 month renewal session) revealed no main effect of CS Type (*p*_*gg*_ = 0.083). *Post-hoc* paired *t*-tests revealed that across renewal phases, similarity was marginally enhanced for CS+CC in comparison to CS-stimuli (t_(22)_ = 2.047, *p =* 0.052, 95% CI [-0.001, 0.155]), but not in comparison to CS+EXT stimuli (*p =* 0.887). (**Fig. 4C**).

**Figure 4.**
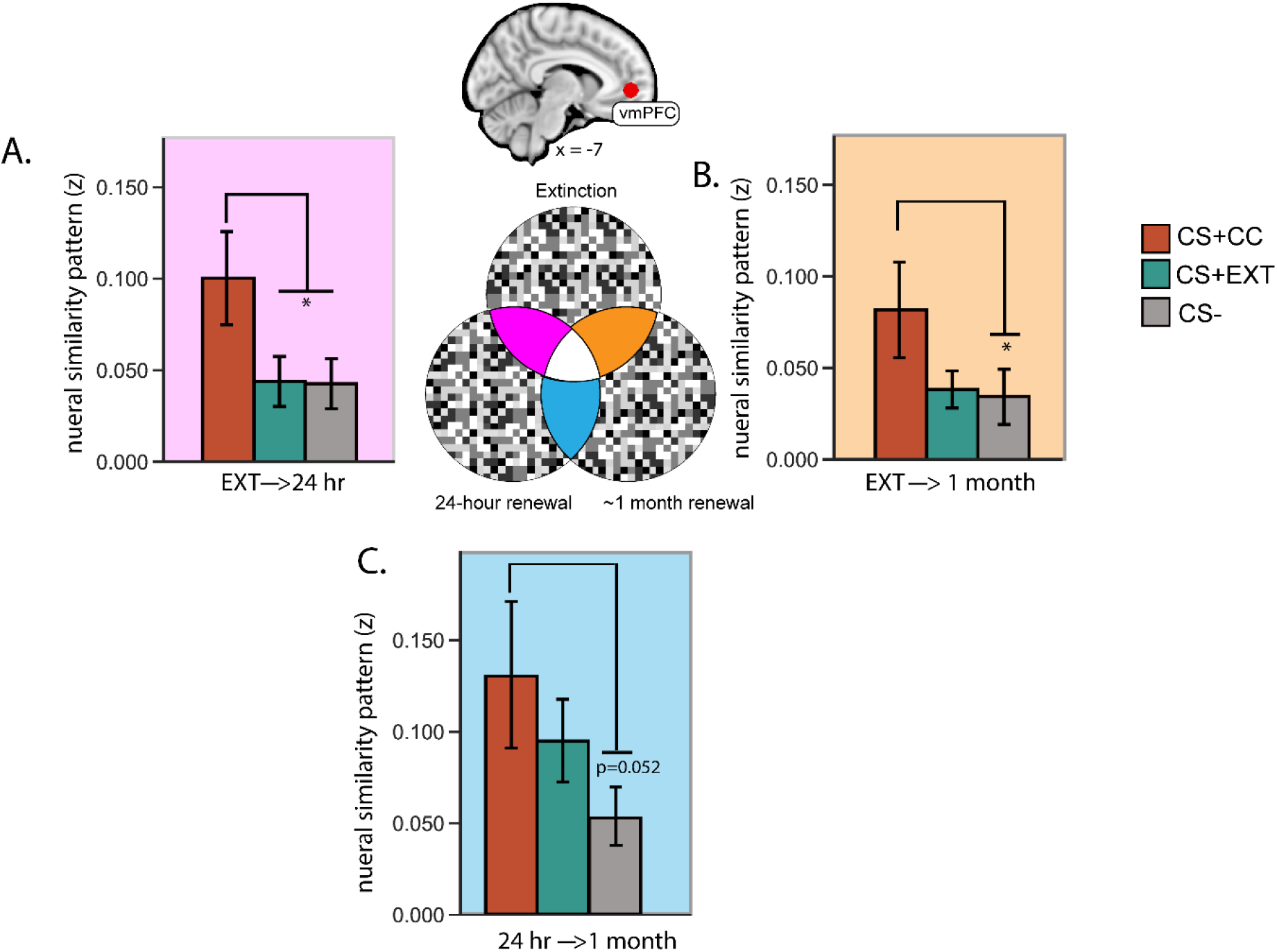
Stimuli that underwent CC were associated with a heightened pattern of similarity in the vmPFC. A.) Pattern similarity from encoding to recent renewal (extinction→24 hour retrieval) in the vmPFC was enhanced for the CS+CC category in comparison to both the CS+EXT and CS-categories. A.) Pattern similarity from encoding to remote renewal (extinction→∼1 month renewal) in the vmPFC was enhanced for the CS+CC category in comparison to the CS-category. C.) Pattern similarity from recent to remote renewal (extinction→24-hour renewal) in the vmPFC was marginally enhanced for the CS+CC category in comparison to the CS-category. Error bars indicate SEM. P<0.001 (***), P<0.01 (**), P<0.05 (*).

## Discussion

As extinction is a transient form of inhibitory learning, there is interest in optimized strategies that more effectively inhibit relapse of extinguished threat. Counterconditioning (CC) may be more effective than standard extinction (Keller et al., 2020), but the neurobehavioral mechanisms of CC in humans have remained unclear. Further, to our knowledge, the long-term neurobehavioral effects of threat attenuation strategies (> 1 week) have remained unexamined in humans. Here we found that, in comparison to standard extinction, rewarded extinction using CC attenuated activity in regions associated with threat appraisal and expression and reduced 24-hour conditioned responses. Twenty-four-hour renewal was accompanied by enhanced functional connectivity between the BLA and NAc for stimuli from the CC category, and connectivity between the vmPFC and CeM for stimuli from the standard extinction category. One-month renewal was associated with reduced conditioned responses and accompanied by connectivity between vmPFC and CeM for both extinction strategies. Representational similarity analysis showed that memory traces of CC are stable in the vmPFC across recent and remote time points.

An overarching question about CC is whether it should simply be considered another form of extinction or whether it operates through different neural mechanisms (Keller et al., 2020). Here, we found that CC attenuated activity in regions associated with threat appraisal and expression (insula, thalamus, dACC, PAG), suggesting that providing a positive experience during extinction may facilitate safety learning. Notably, this finding is consistent with a recent fMRI study in which a shock was replaced with a neutral outcome (a tone) (Dunsmoor et al., 2019). As previously suggested, replacing shock with a non-aversive stimulus might reduce ambiguity and uncertainty otherwise generated when a shock is merely omitted (Dunsmoor et al., 2015).

At 24-hour and 1-month renewal tests, there was a surprising lack of differentiation in whole-brain fMRI activity between the retrieval of CC and standard extinction memories. A more liberal statistical threshold did reveal greater activity for CC in the left amygdala at 24-hour renewal. On one hand this finding may seem counterintuitive, given that the amygdala is critical for threat learning and expression (Phelps and LeDoux, 2005) and conditioned responses were slightly more attenuated by CC. However, the amygdala also responds to rewarding stimuli (Beyeler et al., 2018; Kim et al., 2016; Zhang et al., 2020; Zhang & Li, 2018), and the BLA contains neural populations that code for extinction memory (Herry et al., 2010) and neurons that respond to reward overlap with those involved in extinction (Zhang et al., 2020). Thus, it is possible the amygdala plays an important role in retrieving reward-associations connected with the memory of CC.

We used functional connectivity analysis to further assess the neural differences between CC and standard extinction. At 24-hours, functional connectivity between the vmPFC and CeM was enhanced for standard extinction in comparison to CC; in contrast, functional connectivity between the BLA and the NAc was enhanced for CC in comparison to standard extinction. These findings can be interpreted in the well-explored neurocircuitry of threat extinction in rodents. For instance, infralimbic (rodent homolog of vmPFC) projections to the BLA excite GABAergic intercalated cells that inhibit CeM neurons thereby inhibiting conditioned responses (Amano et al., 2010; Pape & Pare, 2010; Strobel et al., 2015). A BLA to NAc circuit has been identified during rewarded-extinction in rats, and is associated with reduced threat relapse (Correia et al., 2016). Further evidence for the role of the BLA-to-NAc circuit comes from recent studies on rescuing behavioral deficits induced by chronic stress (Dieterich et al., 2021; Sun et al., 2021). Collectively, the present results help extend rodent neurobiological findings to humans and indicate that separate patterns of connectivity dissociate CC from standard extinction. Interestingly, connectivity between vmPFC and CeM was observed at 1-month for both CS types, suggesting that over longer periods of time, extinction recruits medial prefrontal inhibition of the amygdala regardless of the particular threat inhibition strategy. It is worth noting that the 24-hour renewal test served as another standard extinction session, as positive outcomes were not included at test. Thus, the memory of CC at the 1-month test comprised a mix of CC (from Day 1) and standard extinction (from Day 2) that may be reflected in the switch in connectivity from BLA→NAc to vmPFC→CeM over time.

A multivariate RSA was used to further interrogate the fidelity of CC and standard extinction memories. The reactivation of neural activity patterns from extinction were enhanced by CC in the vmPFC both 24-hours and 1-month later. It is notable that the vmPFC showed neural reactivation patterns for CC, as functional connectivity analyses indicated a vmPFC→amygdala connection was selectively enhanced 24-hours following standard extinction but not CC. However, neurobiological evidence shows that activation of the BLA→NAc circuit by rewarded extinction increases activity in the IL to prevent threat relapse (Correia et al., 2016). Thus, CC may likewise enhance involvement of the vmPFC for storing long-term memory traces of safety.

The results from the 1-month retrieval test were intriguing for several reasons. First, although shock expectancy returned slightly, autonomic arousal was remarkably low. This might indicate that both threat attenuation strategies were successful over the long term. It is notable that functional connectivity between the vmPFC and the CeM was evident for both CS+ categories at 1-month (albeit only at a marginal level for CS+EXT), suggesting this is a mechanism for successfully reducing conditioned responses over long durations in humans. It is also important to note that participants were all reportedly free of psychopathology, and thus memory of laboratory conditioned threat might simply weaken over long durations in the healthy brain. This calls for future studies comparing the return of threat over longer intervals in patients with anxiety disorders, particularly posttraumatic stress disorder (PTSD). Threat conditioning is a popular model for PTSD (Mahan & Ressler, 2012) but immediate dysregulated responses to a CS may better reflect Acute Stress Disorder, which refers to the stress symptoms that arise in the first month after a traumatic event (Bryant, 2019). A key criteria in a PTSD diagnosis is the persistence of symptoms at least 1-month following the trauma (American Psychiatric Association, 2013). Importantly, acute stress disorder can develop when PTSD does not, and vice-versa (Bryant, 2010). More research is warranted on the long-term endurance of different extinction strategies in clinical populations who display extinction retrieval deficits.

A limitation of the present study concerns the broad definition of “reward” for the outcomes used to replace shocks in counterconditioning. Simply put, *were the pictures actually rewarding*? More generally, by what operational definition should “reward” be applied? It is worth noting that the pictures used in this study were rated highly in positive valence by a separate group of participants. CC paradigms have employed a wide variety of appetitive outcomes (see Table 1: Keller et al., 2020), as well as different methodology for the subject to obtain the reward (e.g., passively delivered vs. an instrumental behavior, Thomas et al., 2012). From a purely neural perspective, extinction does recruit reward-responsive dopaminergic systems (Kalisch et al., 2019; McNally et al., 2011; Salinas-Hernández and Duvarci, 2021). Further, the mere absence of an expected shock could be construed as a psychological reward (or at least a relief). It is therefore possible that facilitating extinction through any number of strategies simply promotes engagement of a threat-inhibition process that overlaps with reward-responsive neurocircuitry. One way future research could evaluate whether there is a unique effect of “reward”, would be to compare outcomes that vary in reward intensity, such as comparing positive pictures to primary reinforcers, like food or juice, or to compare passive delivery versus instrumental responses (Thomas et al., 2012).

Insofar as Pavlovian extinction serves as a theoretical foundation for exposure therapy, and symptoms frequently return following treatment (Vervliet et al., 2013), examining the neurobehavioral endurance of different threat attenuation strategies is important. These results provide new evidence that the presence of a rewarding stimulus during extinction may boost threat attenuation through an amygdala-striatal pathway, and stabilize memory representations in the vmPFC over long time intervals. These results extend neurobiological findings on the overlap between reward and threat extinction from rodents to healthy humans. While neuroimaging research comparing these strategies in clinical populations is warranted, this type of research could serve as a foundation for translational efforts that result in a paradigm shift for exposure therapy.

